# Top-down Modulation of Brain Responses in Spelling Error Recognition

**DOI:** 10.1101/2023.01.13.523923

**Authors:** Ekaterina Larionova, Zhanna Garakh, Olga Martynova

**Affiliations:** Institute of Higher Nervous Activity and Neurophysiology, Russian Academy of Sciences, Moscow, Russian Federation; Centre for Cognition and Decision Making, Institute for Cognitive Neuroscience, Higher School of Economics, Moscow, Russian Federation

**Keywords:** ERP, top-down control, orthographic decision, reading, visual word recognition, spelling

## Abstract

The task being undertaken can influence orthographic, phonological and semantic processes. In linguistic research, two tasks are most often used: a task requiring a decision in relation to the presented word and a passive reading task which does not require a decision regarding the presented word. The results of studies using these different tasks are not always consistent. This study aimed to explore brain responses associated with the process of recognition of spelling errors, as well as the influence of the task on this process. Event-related potentials (ERPs) were recorded in 40 adults during an orthographic decision task to determine correctly spelled words and words written with errors that did not change the phonology and during the passive reading. During spelling recognition, the early stages up to 100 ms after the stimulus were automatic and did not depend on the requirements of the task. The amplitude of the N1 component (90-160 ms) was greater in the orthographic decision task, but did not depend on the correct spelling of the word. Late word recognition after 350-500 ms was task dependent, but spelling effects were similar across the two tasks: misspelled words evoked an increase in the amplitude of the N400 component related to lexical and semantic processing regardless of the task. In addition, the orthographic decision task modulated spelling effects, this was reflected in an increase in the amplitude of the P2 component (180-260 ms) for correctly spelled words compared with misspelled words. Thus, our results show that spelling recognition involves general lexico-semantic processes independent of the task. Simultaneously, the orthographic decision task modulates the spelling-specific processes necessary to quickly detect conflicts between orthographic and phonological representations of words in memory.

**Highlights:**

- The influence of task on the recognition of correct spelling of words was studied
- Sublexical stages during spelling recognition are task-independent
- The amplitude of the P1, N1 and P600 components is not sensitive to the spelling of words
- The orthographic decision task modulates the spelling processes at 180–260 ms
- Spelling recognition modulates the N400 component regardless of the task

## 1. Introduction

Misspelled words occur in all writing systems. The type of misspelling produced is influenced by the writing system, reading skills, and the spelling tasks (Limpo et al., 2021). However, regardless of the writing system, many misspelled words tend to be phonologically similar to correctly spelled words. Indeed, pseudohomophones, that is, non-words that sound the same as words (for example, the pseudohomophone “brane” for the word “brain” in English), can be a type of natural spelling error. For example, in Russian, all words with errors in the unstressed vowels are pseudohomophones: the pronunciation of the correctly spelled word “peшeниe” (“decision”), in which the first syllable is unstressed, and the misspelled word “pишeниe” (incorrect spelling of the word “peшeниe”) is the same [r^j^I’ʂεn^j^Ije]. Therefore, studies on the recognition of pseudohomophones are closest to the study of spelling errors. Neurophysiological studies of pseudohomophones^1^ sometimes use the silent reading paradigm (Araújo et al., 2015; Kemény et al., 2018; Larionova & Martynova, 2022; Sauseng et al., 2004; Vissers et al., 2006), but more often, they use the lexical decision task wherein the subject needs to determine which stimulus is a word and which is not (Braun et al., 2009; Briesemeister et al., 2009; Costello et al., 2021; González-Garrido et al., 2014, 2015), or similar tasks such as the phonological lexical decision task (Bakos et al., 2018; Hasko et al., 2013), where it is necessary to make a decision about the correspondence of the sound of the stimulus to the word, or the orthographic decision task (Taha & Khateb, 2013), when it is necessary to make a decision on the correct spelling of the word. The spelling decision task is probably the most suitable for studying the processes associated with determining the correct spelling of a word since it gives direct instruction to determine the correct spelling of each stimulus. Nevertheless, the silent reading task is closer to the natural reading process than a lexical decision task since we usually read without making any decision (Chen et al., 2015), which may be why some authors believe that it is better suited for research on visual word recognition (Bermúdez-Margaretto, Beltrán, et al., 2020; Blythe et al., 2020). In addition, the use of such a task makes it possible to understand whether the differentiation of misspelled words and correctly spelled words occurs automatically.

The results obtained in studies with pseudohomophones are not always consistent: in some studies, differences between words and pseudohomophones were observed at about 100-200 ms (ERP components P100, P150, N170) after the presentation of stimuli (Araújo et al., 2015; Braun et al., 2009; González-Garrido et al., 2015; Sauseng et al., 2004), however in others differences were observed much later, in the N400 component and the P600 component following it. It was shown that the amplitude of the N400 component is larger for pseudohomophones than for words (Briesemeister et al., 2009; González-Garrido et al., 2015; Hasko et al., 2013; Vissers et al., 2006), and the P600 component has been shown to both increase (Van de Meerendonk et al., 2011), and decrease (Bakos et al., 2018) in amplitude for pseudohomophones compared to words. This may indicate that spelling recognition can occur both early, at the stages associated with sublexical and lexical processing, and later at the stages of lexico-semantic processing. However, the differences in the research paradigms may be the reason for the conflicting results. In addition, it is still unclear at what stage the task’s requirements affect the processing of verbal information.

The influence of the task on the N400 and P600 components is noted in a number of linguistic studies (Bornkessel-Schlesewsky & Schlesewsky, 2008; Brouwer & Crocker, 2017; Kolk et al., 2003; Schacht et al., 2014). Kolk et al., used two paradigms: in the first task, participants had to identify grammatically incorrect and semantically implausible sentences; in the second task, subjects read these sentences, and after presenting 20% of the sentences answered questions about their content (Kolk et al., 2003). In the first experiment, they found that semantically implausible combinations of words (for example, “the trees that played in the park…”) caused N400/P600 effects, and in the second, only an N400 effect, so the authors conclude that the most common linguistic function P600 is a monitoring or verification function (Kolk et al., 2003). This assumption is consistent with the results of Van de Meerendonk and colleagues (Van de Meerendonk et al., 2011), who showed an increase in the P600 amplitude for spelling violations in words compared to correctly spelled words during passive sentence reading followed by questions about their content, which, according to the authors, reflects reprocessing to check for possible processing errors. Interestingly, in the phonological decision task, the opposite results were obtained (Bakos et al., 2018), which may be due to the fact that in this task, rechecking is required to a lesser extent since pseudohomophones also sound like words.

The N400 component is traditionally associated with lexico-semantic processing, which can be modulated by task conditions (Kiefer & Brendel, 2006; Kutas & Federmeier, 2011; Kutas & Hillyard, 1989). The effect of N400 during the lexical decision task with masked semantic priming was smaller when this task was preceded by a perceptual task, but not when preceded by a semantic task, which, on the contrary, enhanced this effect (Kiefer & Martens, 2010). The authors explained these results using their own model, according to which attentional mechanisms optimize unconscious information processing depending on the goals of the task being performed; that is, unconscious semantic processing was enhanced by the semantic task and weakened by the perceptual task (Kiefer & Martens, 2010). Similar results on the effect of the task on the semantic processing of words and the N400 effect were obtained in another study, while for the early negative wave N200 no effect of the task was found, which, according to the authors, indicates the independence of early automatic semantic processing from task requirements (Marí-Beffa et al., 2005). Brown and colleagues (Brown et al., 2000) showed that in the lexical decision task, the amplitude of N400 is modulated by the proportion of semantically related and unrelated pairs of words, while during silent reading, there is no such modulation; the amplitude of N400 for semantically unrelated target words is greater than the amplitude for related words in both tasks (Brown et al., 2000). According to some authors, this may indicate the existence of various priming mechanisms: automatic priming and controlled priming, which are associated with expectation (Brown et al., 2000; Silva-Pereyra et al., 2003). Accordingly, the N400 component, according to some authors, is sensitive to both automatic and controlled processing components (Kiefer & Brendel, 2006; Kutas & Federmeier, 2011; Kutas & Hillyard, 1989). Meade and colleagues (Meade et al., 2019) found orthographic neighborhood effects associated with the N400 component for words in both the lexical decision task and the letter search task, but the N400 neighborhood effect for pseudowords has only been found in the lexical decision task. The authors suggested that the neighborhood effect for words is due to automatic word recognition processes, while the neighborhood effect for pseudowords is associated with a task-specific process of enhancing global lexical activity for making a lexical decision (Meade et al., 2019). Unfortunately, a comparison of the modulation of the N400 component for pseudohomophones in different paradigms has not been made before. However, in general, an increase in the amplitude of the N400 for pseudohomophones than for words is most commonly found in studies (e.g., Briesemeister et al., 2009; González-Garrido et al., 2015; Hasko et al., 2013; Larionova & Martynova, 2022; Vissers et al., 2006), suggesting that it is independent of the paradigm used.

There is evidence that a task can modulate not only the lexico-semantic but also the phonological processes necessary to perform a specific task (e.g., Barnea & Breznitz, 1998; Chen et al., 2015). For example, Barnea and Breznitz (Barnea & Breznitz, 1998) found an increase in the latency and amplitude of the P200 and N400 components during a phonological task (rhyme-nonrhyme decision task) compared to a spelling task (orthographic similarity decision task). Chen and colleagues (Chen et al., 2013) using EEG/MEG and fMRI, found a task effect 150 ms after word presentation, greater activation of the left precentral areas was observed during the silent reading task than during lexical decision tasks, which the authors attribute to increased phonological activation during silent reading compared to the lexical decision task. In addition, depending on the task, the topography of the components may also differ: N350 was highest in the left frontal area in the phonological task (to assess whether pairs of words rhyme), in contrast to the semantic task (to assess whether pairs of words are semantically related), in which this component had a two-sided distribution, while N150 did not differ between these tasks (Spironelli & Angrilli, 2007).

Despite the traditional view that the early stages of word recognition are automatic and independent of task requirements (Herdman & Takai, 2013), task-related top-down processes have been shown to influence the early stages of verbal information processing at 250 ms after the stimulus (Chen et al., 2015; Faísca et al., 2019; Strijkers et al., 2015; F. Wang & Maurer, 2017). During the implicit reading task, in which subjects had to press a button when stimuli were repeated and there was no instruction to read words, N1 had a larger amplitude for correctly spelled word stimuli than for pseudowords, but during reading aloud this modulation of N1 was absent – the amplitude for words and pseudowords did not differ (Faísca et al., 2019). Strikers and colleagues showed that the word frequency effect appeared as early as 120 ms after stimulus presentation in the semantic categorization task but only 220 ms after stimulus presentation in the font color categorization task, where linguistic processing is not required (Strijkers et al., 2015). In addition, other work showed that more pronounced early frequency effects (an increase in N170 amplitude) were observed in the lexical decision task than during silent reading (Chen et al., 2015). Thus, the task may also influence early lexical effects.

Task modulation of brain responses in spelling error recognition has not previously been studied. At the same time, data from neurophysiological studies on the ERP components associated with pseudohomophone processing remain contradictory, which may be due to a task influence. There are studies showing that the task can influence orthographic, phonological, and semantic processes. The process of spelling recognition may be associated with all these processes. The purpose of this work is to study top-down modulation of the dynamics of recognition of correctly spelled words and misspelled words. For this, we analyzed the ERP components associated with various stages of word processing. As stimuli, we used words with real errors in the Russian language, which are pseudohomophones. We hypothesize that an orthographic decision task, in which the subject is required to determine the correct spelling of a word, influences the processes associated with recognition of the correct spelling and enhances spelling effects compared to the passive reading task, this should be reflected in earlier differences in brain responses. If the recognition of spelling errors occurs automatically, then the effects associated with recognition of correct spelling will be the same regardless of the task.

## 2. Methods

### 2.1. Participants

Forty healthy volunteers participated in the study (25 women and 15 men). The subjects were right-handed 20 to 35 year old native Russian speakers without speech disorders, neurological diseases, or psychiatric diseases, and had normal or corrected to normal vision. The average age of the subjects was 26.2 (SD 5.2) years; the average level of education was 14.7 (SD 1.6) years in education. The study was carried out in compliance with the principles of the Declaration of Helsinki, the study was approved by the Ethics Committee, before the start of the experiment, all subjects signed informed consent to participate in the study.

### 2.2. Stimuli

All stimuli were nouns spelled with or without orthographic errors, that consisted of 5-7 letters; only one error was allowed in misspelled words. There were two sets of stimuli. Examples and characteristics for each set of stimuli are presented in Table 1. It is important to note that the misspelled words sound the same as correctly spelled words; that is, they are pseudohomophones. The stimuli in each set were composed in such a way that among them there were words of different frequencies of occurrence, as determined by a word frequency dictionary (Lyashevskaya & Sharov, 2009). Word frequency, word length, and the orthographic neighborhood size did not differ significantly for the two categories of stimuli and between sets (t-test, p > 0.05). Spelling errors were predominantly in the first syllable: for the Set 1, there were 8 words in which the error was in the second syllable; for the Set 2, there were 13 words with an error in the second syllable. One type of spelling error was used – errors in unstressed vowels, as this is a common type of error in Russian. Correctly spelled words also contained an unstressed vowel, but it was spelled correctly.

**Table 1.**
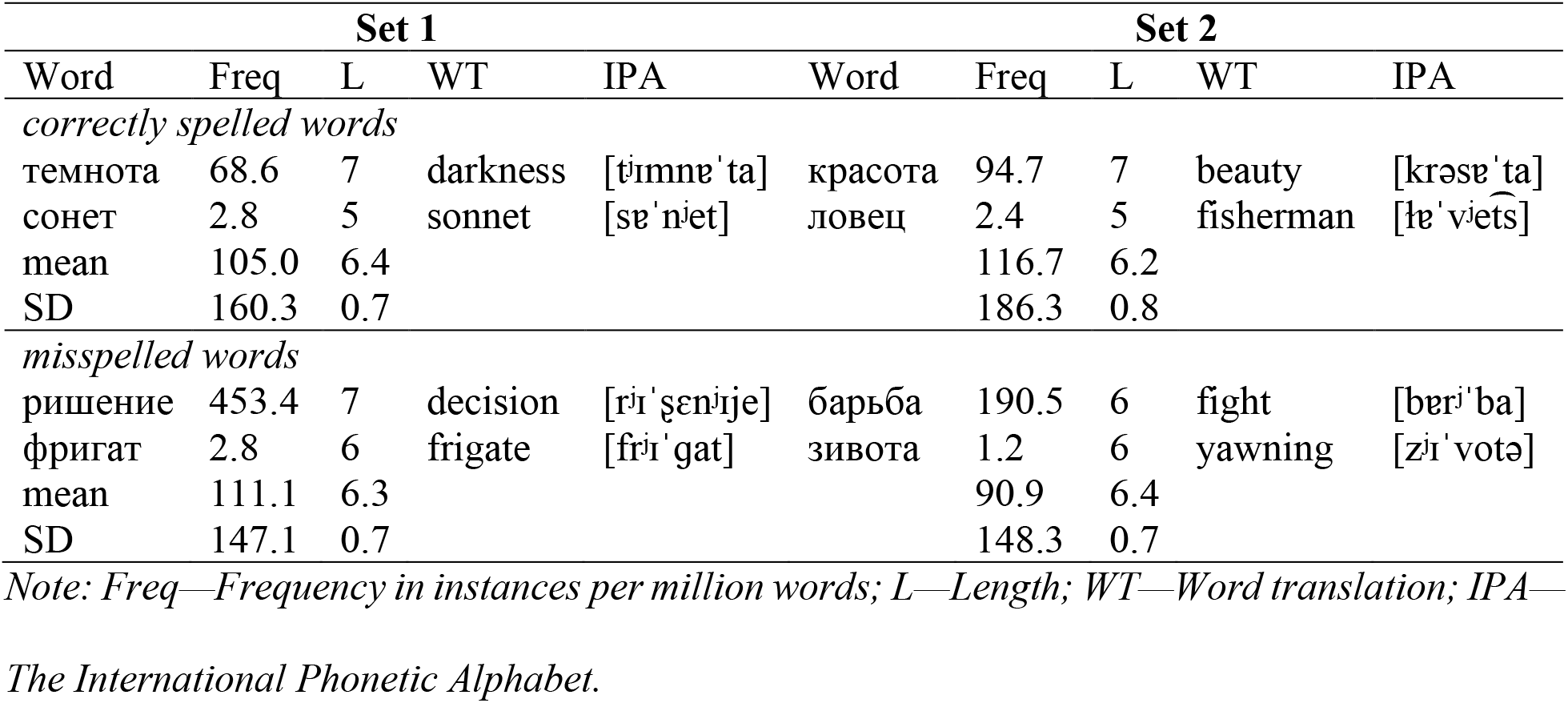
Examples and characteristics of stimuli.

### 2.3. Procedure

During the experiment, participants sat in a darkened room in a comfortable chair at a distance of about 1 m from the monitor. Stimuli were presented in random order on a 19-inch LG FLATRON L1952T monitor using the PsychoPy Experiment Builder v3.0.7 software (Peirce et al., 2019). The stimuli were written in white Liberation Sans on a black background, font size 125 pt.

The subjects had to complete two tasks (the passive reading task and the orthographic decision task); the total duration of the tasks did not exceed 20 minutes, including short breaks. The order of tasks was fixed: the reading task, then the orthographic decision task. This task order was chosen to avoid intentional spelling recognition in a reading task in which subjects were only asked to read words silently. Each task used a different but well-controlled set of stimuli because processing the same stimuli twice could induce a bias and affect the associated neural responses. Two sets of stimuli alternated between tasks: 26 participants received set 1 in the reading task and set 2 in the orthographic decision task; 14 participants received set 2 in the reading task and set 1 in the orthographic decision task.

In the passive reading task, the subjects were asked to silently read the words presented on the screen: 47 words misspelled in an unstressed vowel and 46 correctly spelled words with an unstressed vowel. The presentation time was 300 ms; the interstimulus interval varied randomly from 1500 to 2200 ms.

In the orthographic decision task, the subjects were asked to determine whether there is an error in the word or not. This task required a motor response. The motor response was performed using a Logitech F310 gamepad by pressing the left and right shoulder buttons. The instruction for one half of the participants was as follows: “If there is a mistake in the word, press the right button. If there is no mistake in the word, press the left button.” The instruction for the other half of the participants was as follows: “If there is a mistake in the word, press the left button. If there is no mistake in the word, press the right button.” The number of stimuli was the same as in the first task: 47 words misspelled in an unstressed vowel and 46 correctly spelled words with an unstressed vowel. The presentation time was 300 ms; the interstimulus interval was counted from the motor response and varied randomly from 1500 to 2200 ms. If there was no response within 5000 ms, the next stimulus was presented.

### 2.4. EEG data acquisition and analysis

EEG was recorded using the Encephalan amplifier (Medicom, Russia) from 19 electrodes: Fp1, Fp2, F3, F4, F7, F8, C3, C4, T3, T4, T5, T6, P3, P4, O1, O2, Fz, Cz, Pz, arranged in a 10-20 system. The monopolar ipsilateral design of EEG recording was used with reference electrodes placed on the mastoids. The sampling rate was 250 Hz. The impedance did not exceed 10 kΩ. The event-related potentials were processed using the Brain Vision Analyzer 2.0.4 software (Brain Products, GmbH, Munich, Germany), including 0.5–30 Hz pre-filtering and removal of artifacts using independent component analysis. ERPs were averaged separately for each type of stimulus in each task within a time window of 300 ms before stimulus presentation and 1500 ms after. The incorrect trials in the orthographic decision task were not discarded from the ERP analysis: firstly, in this way, we equalized the number of averagings in the two compared tasks, and secondly, in order to equalize the experimental conditions, in the reading task we did not control task performance. The minimum number of stimuli for averaging was 40 words.

To determine the time intervals for the analysis of the main components of the ERPs in the two tasks, we calculated the global field power (Global Field Power, Figure 1), which allowed us to choose the optimal periods of stable topography (Lehmann & Skrandies, 1980). The main peaks were observed in the intervals 70-90 ms (approximately corresponding to the parietal-occipital-temporal component of P1), 90-160 ms (approximately corresponding to the parietal-occipital-temporal component of N1), 180-260 ms (approximately corresponding to the P2 component, which had a wide topography), 350-750 ms (approximately corresponding to the N400 component, as well as the P600 following it, so this interval was divided into two – 350-500 ms and 500-750 ms) (Figure 1).

**Figure 1.**
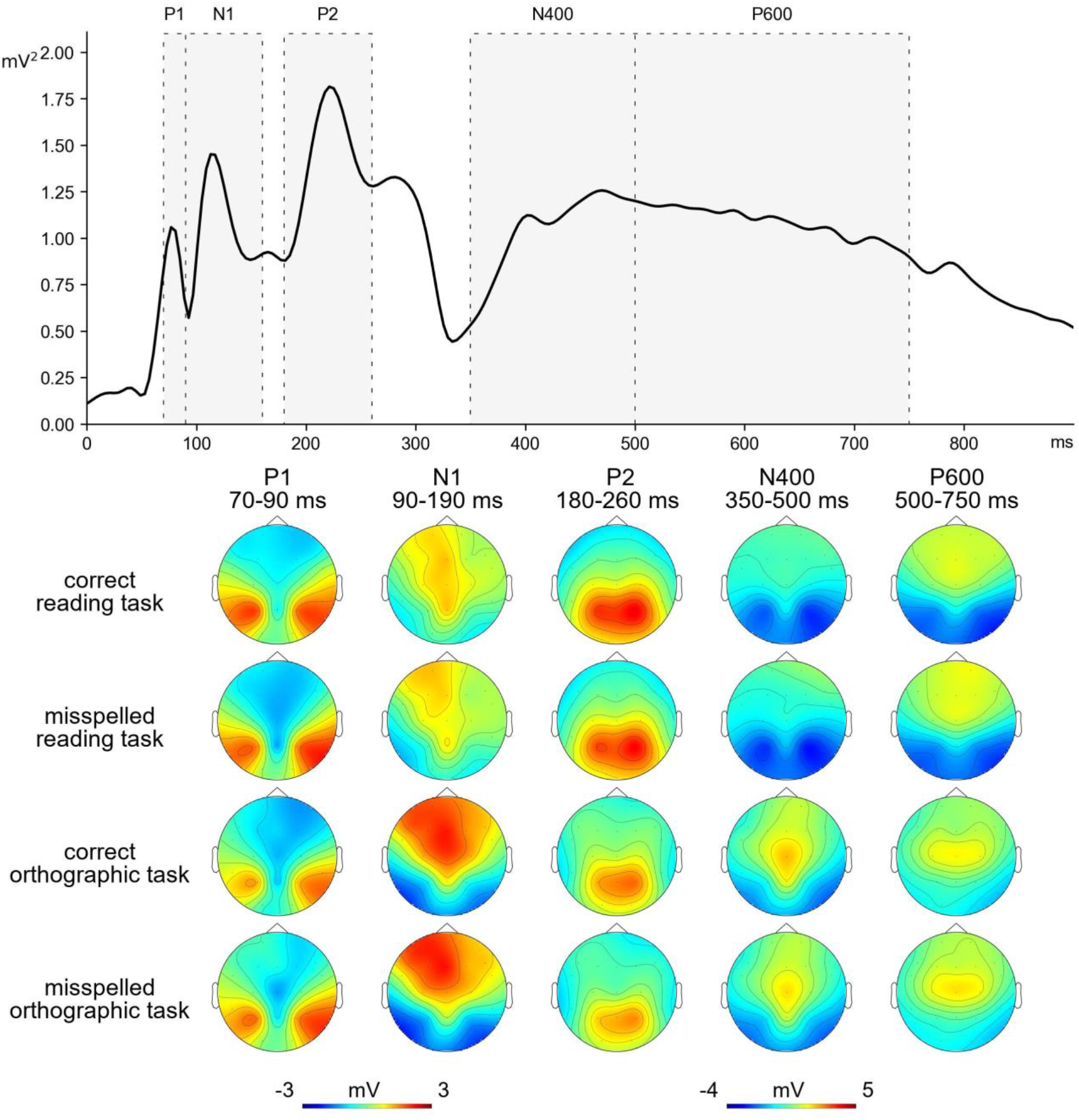
Top: Global field power (GFP, all electrodes) averaged across tasks and experimental conditions with indication of the studied time intervals. Bottom: Topographic maps of the P1, N1, P2, N400, and P600 components.

Statistical analysis of the data was carried out using the STATISTICA 8 software (Statsoft, Tulsa, OK, USA). Analysis of variance with repeated measures (RM ANOVA) was applied to the mean ERP amplitude for each region of interest in each time window. The electrodes were divided into 9 regions of interest: left anterior region (Fp1, F3, F7), middle anterior region (Fz), right anterior region (Fp2, F4, F8), left central region (T3, C3), central region (Cz), right central region (T4, C4), left posterior region (T5, P3, O1), posterior central region (Pz), and right posterior region (T6, P4, O2). We averaged the ERP amplitude across the electrodes in each region of interest. The analysis of variance included the following factors: task (2 levels: passive reading and orthographic decision task), spelling (2 levels: correctly spelled words and misspelled words), laterality (3 levels: left, right, and middle line of electrodes) and electrode position (3 levels: anterior, central, and posterior). In addition, we used a Set of stimuli (Set 1 and Set 2) as a factor in the statistical model. If the observed effect interacted with the Set factor, this effect was not discussed further.

Since the P1 and N1 components, in contrast to the P2, N400 and P600 components with a wide topography, are localized in the parietooccipitotemporal regions (Figure 1), only two regions of interest were included in the analysis of variance for these components: the left and right posterior regions, similarly to other studies that have examined these components (e.g., Faísca et al., 2019; Maurer et al., 2005). We were interested in the effects of task and spelling, as well as their interaction with other factors. When determining the reliability of the effects, the Greenhouse-Geiser correction was taken into account. The Bonferroni correction was used for post hoc analysis. Partial eta squared (η^2^_p_) was applied as a measure of effect size.

## 3. Results

The topography of all studied ERP components, with the exception of the P600 wave, were similar in the two tasks (Figure 1). Examples of average ERPs are shown in Figure 2.

**Figure 2.**
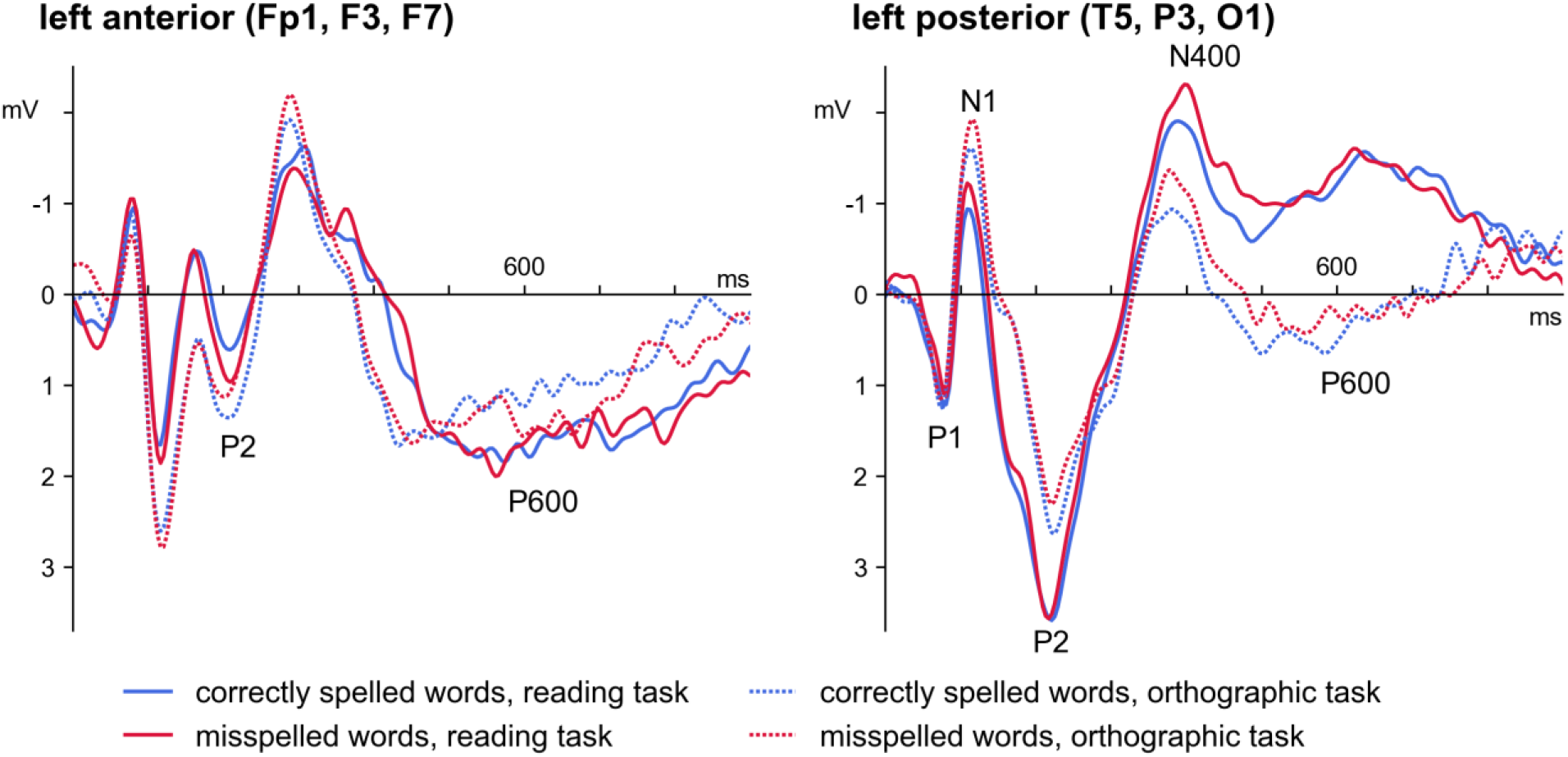
Grand average ERPs in the left anterior and left posterior regions of interest.

The results are presented in Tables 2 and 3.

**Table 2.** Summary ANOVA for P1 and N1.

|  | 70–90 ms (P1) | 90–160 ms (N1) |
| --- | --- | --- |
| Task | $F(1, 38) = 0.25, p = 0.62$ | **$F(1, 38) = 30.10, p = 0.000003$** |
| Task × Set | $F(1, 38) = 0.26, p = 0.61$ | $F(1, 38) = 0.21, p = 0.65$ |
| Spelling | $F(1, 38) = 0.68, p = 0.41$ | $F(1, 38) = 3.63, p = 0.07$ |
| Spelling × Set | $F(1, 38) = 2.20, p = 0.16$ | $F(1, 38) = 0.09, p = 0.77$ |
| Task × Spelling | $F(1, 38) = 0.18, p = 0.67$ | $F(1, 38) = 0.90, p = 0.35$ |
| Task × Spelling × Set | $F(1, 38) = 1.35, p = 0.25$ | **$F(1, 38) = 6.28, p = 0.02$** |
| Task × Laterality | $F(1, 38) = 0.18, p = 0.67$ | $F(1, 38) = 0.22, p = 0.64$ |
| Task × Laterality × Set | $F(1, 38) = 0.29, p = 0.59$ | $F(1, 38) = 0.02, p = 0.88$ |
| Spelling × Laterality × Set | $F(1, 38) = 0.21, p = 0.65$ | $F(1, 38) = 0.45, p = 0.51$ |
| Spelling × Laterality | $F(1, 38) = 0.08, p = 0.78$ | $F(1, 38) = 0.03, p = 0.86$ |
| Task × Spelling × Laterality | $F(1, 38) = 0.90, p = 0.35$ | $F(1, 38) = 1.52, p = 0.23$ |
| Task × Spelling × Laterality × Set | $F(1, 38) = 0.30, p = 0.59$ | $F(1, 38) = 0.04, p = 0.85$ |

**Table 3.**
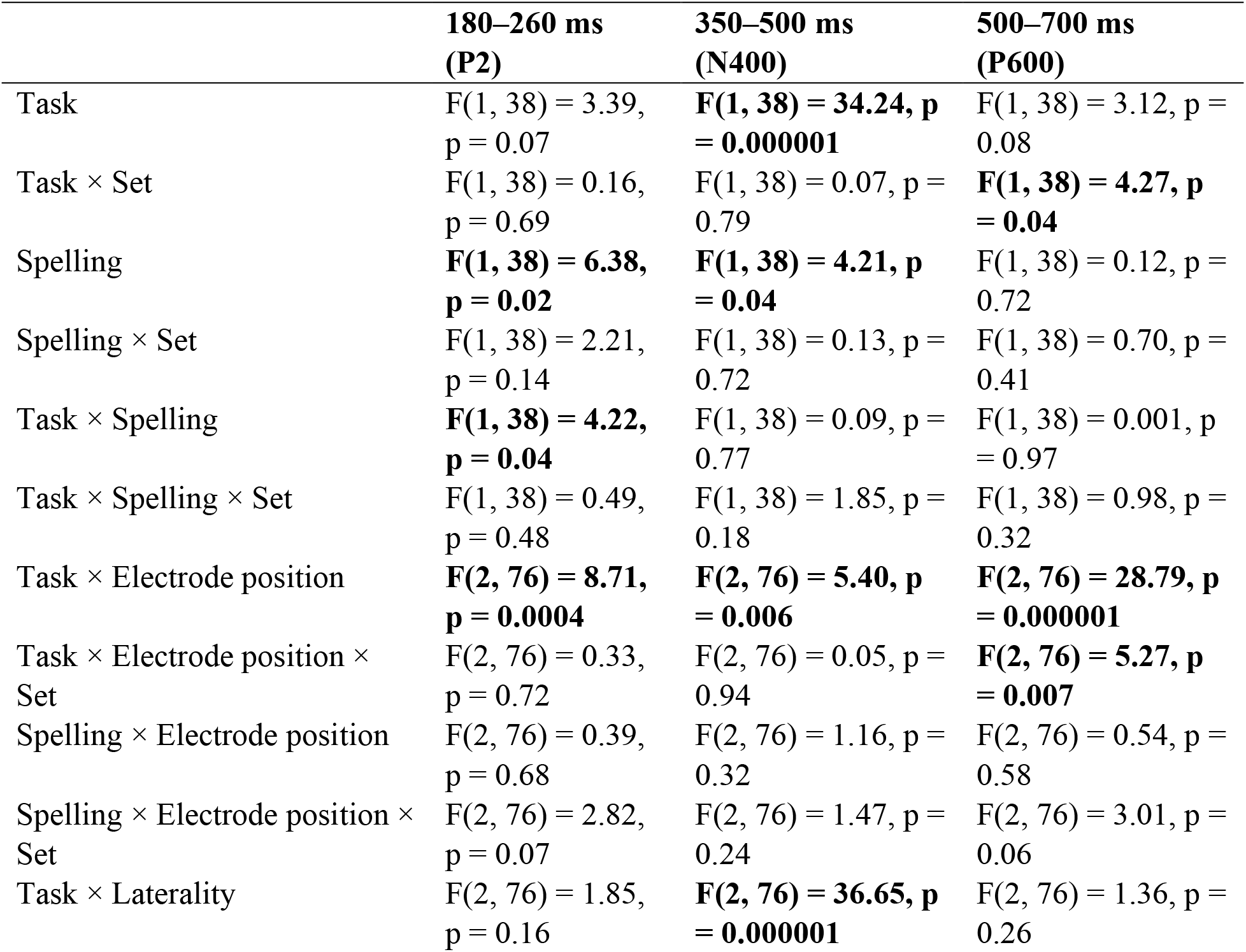
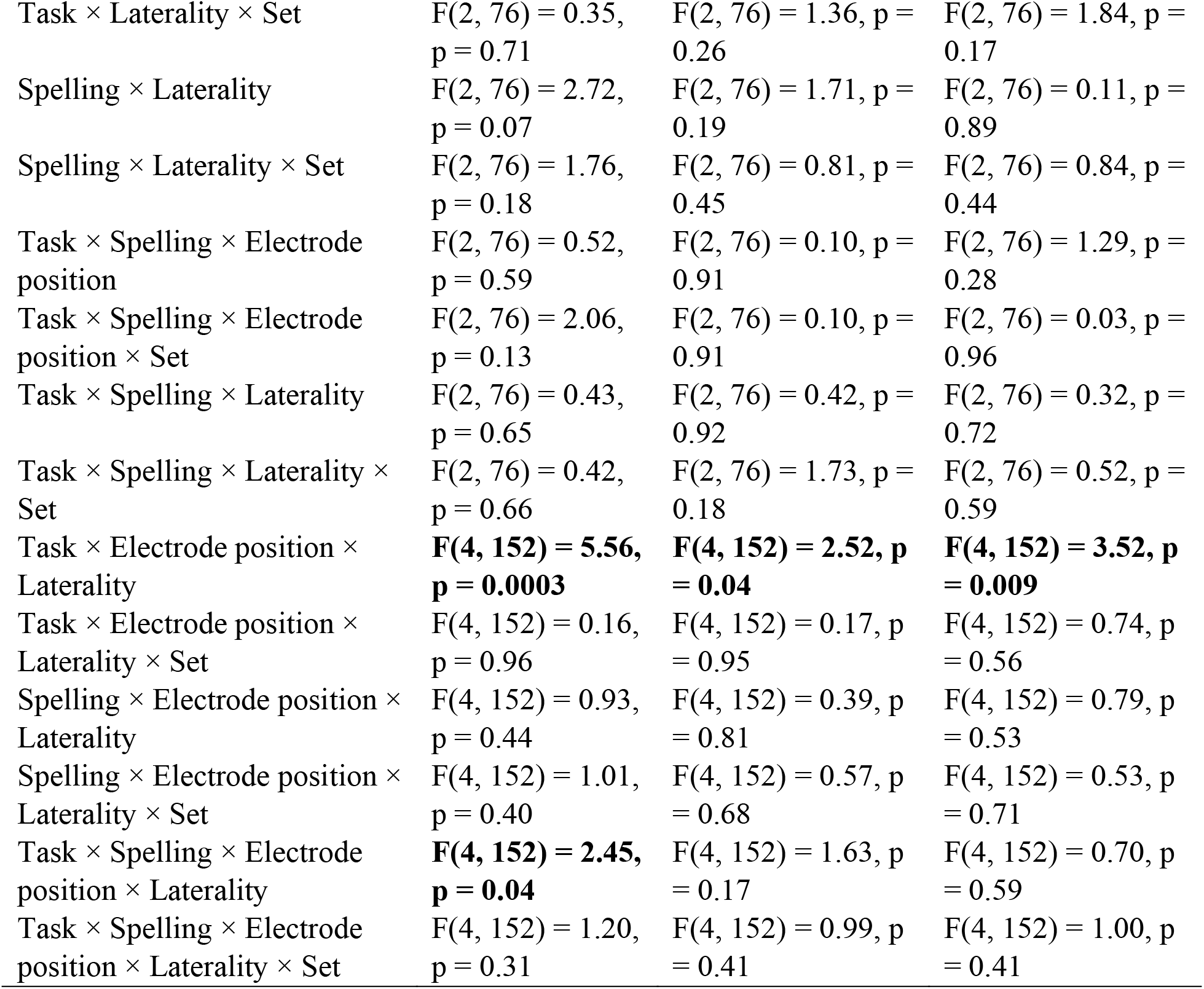
Summary ANOVA for P2, N400 and P600.

|  | **180–260 ms**<br>**(P2)** | **350–500 ms**<br>**(N400)** | **500–700 ms**<br>**(P600)** |
| --- | --- | --- | --- |
| Task | $F(1, 38) = 3.39$,<br>$p = 0.07$ | **$F(1, 38) = 34.24$, $p = 0.000001$** | $F(1, 38) = 3.12$, $p = 0.08$ |
| Task × Set | $F(1, 38) = 0.16$,<br>$p = 0.69$ | $F(1, 38) = 0.07$, $p = 0.79$ | **$F(1, 38) = 4.27$, $p = 0.04$** |
| Spelling | **$F(1, 38) = 6.38$,**<br>**$p = 0.02$** | **$F(1, 38) = 4.21$, $p = 0.04$** | $F(1, 38) = 0.12$, $p = 0.72$ |
| Spelling × Set | $F(1, 38) = 2.21$,<br>$p = 0.14$ | $F(1, 38) = 0.13$, $p = 0.72$ | $F(1, 38) = 0.70$, $p = 0.41$ |
| Task × Spelling | **$F(1, 38) = 4.22$,**<br>**$p = 0.04$** | $F(1, 38) = 0.09$, $p = 0.77$ | $F(1, 38) = 0.001$, $p = 0.97$ |
| Task × Spelling × Set | $F(1, 38) = 0.49$,<br>$p = 0.48$ | $F(1, 38) = 1.85$, $p = 0.18$ | $F(1, 38) = 0.98$, $p = 0.32$ |
| Task × Electrode position | **$F(2, 76) = 8.71$,**<br>**$p = 0.0004$** | **$F(2, 76) = 5.40$, $p = 0.006$** | **$F(2, 76) = 28.79$, $p = 0.000001$** |
| Task × Electrode position × Set | $F(2, 76) = 0.33$,<br>$p = 0.72$ | $F(2, 76) = 0.05$, $p = 0.94$ | **$F(2, 76) = 5.27$, $p = 0.007$** |
| Spelling × Electrode position | $F(2, 76) = 0.39$,<br>$p = 0.68$ | $F(2, 76) = 1.16$, $p = 0.32$ | $F(2, 76) = 0.54$, $p = 0.58$ |
| Spelling × Electrode position × Set | $F(2, 76) = 2.82$,<br>$p = 0.07$ | $F(2, 76) = 1.47$, $p = 0.24$ | $F(2, 76) = 3.01$, $p = 0.06$ |
| Task × Laterality | $F(2, 76) = 1.85$,<br>$p = 0.16$ | **$F(2, 76) = 36.65$, $p = 0.000001$** | $F(2, 76) = 1.36$, $p = 0.26$ |
| Task × Laterality × Set | F(2, 76) = 0.35,<br>p = 0.71 | F(2, 76) = 1.36, p =<br>0.26 | F(2, 76) = 1.84, p =<br>0.17 |
| Spelling × Laterality | F(2, 76) = 2.72,<br>p = 0.07 | F(2, 76) = 1.71, p =<br>0.19 | F(2, 76) = 0.11, p =<br>0.89 |
| Spelling × Laterality × Set | F(2, 76) = 1.76,<br>p = 0.18 | F(2, 76) = 0.81, p =<br>0.45 | F(2, 76) = 0.84, p =<br>0.44 |
| Task × Spelling × Electrode<br>position | F(2, 76) = 0.52,<br>p = 0.59 | F(2, 76) = 0.10, p =<br>0.91 | F(2, 76) = 1.29, p =<br>0.28 |
| Task × Spelling × Electrode<br>position × Set | F(2, 76) = 2.06,<br>p = 0.13 | F(2, 76) = 0.10, p =<br>0.91 | F(2, 76) = 0.03, p =<br>0.96 |
| Task × Spelling × Laterality | F(2, 76) = 0.43,<br>p = 0.65 | F(2, 76) = 0.42, p =<br>0.92 | F(2, 76) = 0.32, p =<br>0.72 |
| Task × Spelling × Laterality ×<br>Set | F(2, 76) = 0.42,<br>p = 0.66 | F(2, 76) = 1.73, p =<br>0.18 | F(2, 76) = 0.52, p =<br>0.59 |
| Task × Electrode position ×<br>Laterality | **F(4, 152) = 5.56,**<br>**p = 0.0003** | **F(4, 152) = 2.52, p =**<br>**0.04** | **F(4, 152) = 3.52, p =**<br>**0.009** |
| Task × Electrode position ×<br>Laterality × Set | F(4, 152) = 0.16,<br>p = 0.96 | F(4, 152) = 0.17, p =<br>0.95 | F(4, 152) = 0.74, p =<br>0.56 |
| Spelling × Electrode position ×<br>Laterality | F(4, 152) = 0.93,<br>p = 0.44 | F(4, 152) = 0.39, p =<br>0.81 | F(4, 152) = 0.79, p =<br>0.53 |
| Spelling × Electrode position ×<br>Laterality × Set | F(4, 152) = 1.01,<br>p = 0.40 | F(4, 152) = 0.57, p =<br>0.68 | F(4, 152) = 0.53, p =<br>0.71 |
| Task × Spelling × Electrode<br>position × Laterality | **F(4, 152) = 2.45,**<br>**p = 0.04** | F(4, 152) = 1.63, p =<br>0.17 | F(4, 152) = 0.70, p =<br>0.59 |
| Task × Spelling × Electrode<br>position × Laterality × Set | F(4, 152) = 1.20,<br>p = 0.31 | F(4, 152) = 0.99, p =<br>0.41 | F(4, 152) = 1.00, p =<br>0.41 |

No significant effects were found for the 70–90 ms (P1) time window.

The main effect of Task was significant for the 90–160 ms (N1) time window F(1, 38) = 30.10, p = 0.000003, η^2^_p_ = 0.44: the N1 amplitude was more negative in the orthographic decision task than in the passive reading task.

The main effect of Spelling was significant for the 180-260 ms (P2) time window F(1, 38) = 6.38, p = 0.02, η^2^_p_ = 0.14: the P2 amplitude was more positive for correctly spelled words than for misspelled words. In addition, we observed significant interaction of the factors Task × Spelling: F(1, 38) = 4.22, p = 0.04, η^2^_p_ = 0.10. The post-hoc analysis showed that the P2 amplitude was more positive in the reading task compared to the orthographic decision task for only misspelled words (p = 0.0003) and also that the amplitude for correctly spelled words was greater than for misspelled words only in the orthographic task (p = 0.0001). In the same time window, we observed significant interaction of the factors Task × Electrode position: F(2, 76) = 8.71, p = 0.0004, η^2^_p_ = 0.19. The post-hoc analysis showed that the P2 amplitude was larger in the reading task compared with the orthographic decision task in the parietal-temporal-occipital region (T5, P3, O1, Pz, T6, P4, O2, p = 0.0002). In addition, we revealed significant interaction of the factors Task × Electrode position × Laterality: F(4, 152) = 5.56, p = 0.0003, η^2^_p_ = 0.13. The post-hoc analysis showed that the ERP amplitude in the 180-260 ms time window in the left frontal region (Fp1/F3/F7, p = 0.04) was greater in the orthographic task than in the reading task; in the central (Cz, p < 0.000001), right central (T4, C4, p = 0.02), left posterior (T5, P3, O1, p < 0.000001), posterior central (Pz, p < 0.000001), and right posterior (T6, P4, O2, p < 0.000001) areas ERP amplitudes were larger during reading than during the orthographic task. And finally, we observed four-way interaction of the factors Task × Spelling × Electrode position × Laterality: F(4, 152) = 2.45, p = 0.04, η^2^_p_ = 0.07. Further analysis in each area of interest showed a significant interaction of Task × Spelling (F(1, 39) = 4.63, p = 0.04, η^2^_p_ = 0.11) in the left frontal region. The post-hoc analysis showed that the amplitude for correctly spelled words was greater in the orthographic task than in the reading task (p = 0.003). The main effect of Spelling (F(1, 39) = 4.63, p = 0.02, η^2^_p_ = 0.14) was found in the left central region: the amplitude for correctly spelled words was greater than for misspelled words. In the central region, effects of Task (F(1, 39) = 7.16, p = 0.01, η^2^_p_ = 0.16, more positive P2 amplitude for reading compared to the orthographic task) and Spelling (F(1, 39) = 4.31, p = 0.04, η^2^_p_ = 0.10, more positive P2 amplitude for correctly spelled words compared to misspelled words) and interaction of Task × Spelling (F(1, 39) = 4.64, p = 0.04, η^2^_p_ = 0.11) were discovered. The post-hoc analysis showed that the P2 amplitude in the central region was more positive in the reading task compared to the orthographic decision task for only misspelled words (p = 0.0001) and also that the amplitude for correctly spelled words was greater than for misspelled words only in the orthographic task (p = 0.0001). The main effect of Task (F(1, 39) = 32.78, p = 0.000001, η^2^_p_ = 0.46) was found in the left posterior region: the P2 amplitude was greater during reading compared than during the orthographic task. In the posterior central region, effects of Task (F(1, 39) = 6.39, p = 0.01, η^2^_p_ = 0.14, more positive P2 amplitude for reading compared to orthographic task) and Spelling (F(1, 39) = 4.16, p = 0.04, η^2^_p_ = 0.10, more positive P2 amplitude for correctly spelled words compared to misspelled words) were discovered. The main effect of Task (F(1, 39) = 37.01, p = 0.000001, η^2^_p_ = 0.49) was also observed in the right posterior region, where post hoc analysis showed that the P2 amplitude was greater during reading than during the orthographic decision task.

In the 350-500 ms (N400) time window, the main effect of Task (F(1, 38) = 34.24, p = 0.000001, η^2^_p_ = 0.47) was significant: a more negative wave was observed in the reading task than in the orthographic decision task. In addition, we identified the main effect of Spelling (F(1, 38) = 4.21, p = 0.04, η^2^_p_ = 0.10): a more negative wave was observed for misspelled words than for correctly spelled words. In the same time window, we observed significant interaction of the factors Task × Electrode position: F(2, 76) = 5.40, p = 0.006, η^2^_p_ = 0.12. The post-hoc analysis showed that the N400 amplitude was more negative in the reading task compared with the orthographic decision task in the anterior region (Fp1, F3, F7, Fz, Fp2, F4, F8, p < 0.000001), in the central region (T3, C3, Cz, T4, C4, p < 0.000001) and in the posterior region (T5, P3, O1, Pz, T6, P4, O2, p < 0.000001). For significant interaction of Task × Laterality (F(2, 76) = 36.65, p = 0.000001, η^2^_p_ = 0.49), post-hoc analysis showed that a more negative wave was observed in the reading task than in the orthographic decision task in the left (Fp1, F3, F7, T3, C3, T5, P3, O1, p < 0.000001), central (Fz, Cz, Pz, p < 0.000001), and right (Fp2, F4, F8, T4, C4, T6, P4, O2, p < 0.000001) electrodes. And finally, we observed a significant interaction of Task × Electrode position × Laterality: F(4, 152) = 2.52, p = 0.04, η^2^_p_ = 0.06. The post-hoc analysis showed that a more negative wave was observed in the reading task than in the orthographic decision task in all leads (p < 0.000001).

In the 500-750 ms (P600) time window, significant interaction of Task × Electrode position (F(2, 76) = 28.79, p = 0.000001, η^2^_p_ = 0.43) and significant interaction of Task × Electrode position × Laterality (F(4, 152) = 3.52, p = 0.009, η^2^_p_ = 0.18) were discovered. The post-hoc analysis showed that the amplitude in the left anterior region (Fp1/F3/F7, p = 0.002) and middle anterior region (Fz, p = 0.0004) was more positive in the reading task than in the orthographic decision task, but in the central (T3/C3, p = 0.0003; Cz, p < 0.000001; T4/C4, p = 0.001) and posterior (T5/P3/O1, p < 0.000001; Pz, p < 0.000001; T6/P4/O2, p < 0.000001) areas was more positive in the orthographic decision task than in the reading task.

## 4. Discussion

Prior ERP experiments have shown that task may modulate brain responses and some processes in word recognition. In the present study, we investigated the influence of task on the ERP components associated with the recognition of the correctness of spelled words. ERP results show an influence of the task on almost all stages of the processing of verbal information. The orthographic decision task modulated the spelling effects in a 180-260 ms time window, corresponding to the P2 component. First, we observed an increase in the amplitude of the P2 component in the left frontal area for correctly spelled words in the orthographic task versus the reading task. Second, we showed an increase in the amplitude of the P2 component in the central region on correctly spelled words over words written with an error.

At the earliest time epoch of analysis (70–90 ms), related to P1, we found no effect of the task on brain responses. The P1 component is associated with the analysis of the physical parameters of the stimulus, but it can also be connected with early attentional processes (Araújo et al., 2015; Klimesch, 2011; Proverbio & Zani, 2003). In addition, this stage is associated with sublexical processes and ERP studies have shown that the P1 component of ERPs was larger for atypical letter combinations than typical ones when the physical parameters of the stimuli did not differ (Araújo et al., 2015; Coch & Mitra, 2010; Hauk et al., 2006). In our study, the physical parameters of the stimuli (color, size, orientation, screen position) were the same in the two tasks, and the amplitude of the P1 component did not differ during the reading task nor during the orthographic decision task. Similar results have been obtained in other studies. For instance, when comparing a task in which subjects had to pay attention to the repetition of verbal stimuli and the task of reading aloud, the effect of the task for the P1 component was absent, but was observed for the later N1 (Faísca et al., 2019). When comparing word and pseudoword reading aloud and the lexical decision task for the same stimuli, no differences were found in the topography and amplitude of the P1 component, so the authors conclude that very early low-level visual processes are common to both tasks (Mahé et al., 2015). When processing correctly spelled words and misspelled words, the early sublexical stages up to 100 ms are probably automatic and do not depend on the task’s requirements.

For the other early wave (90–160 ms), our results indicate a significant task effect: the negative N1 wave was greater in the orthographic decision task than in the reading task. On one hand, this result corroborates the findings of a great deal of previous work showing that the amplitude of the N170 (or N1) component can be modulated by selective attention (e.g., Aranda et al., 2010; Hillyard & Anllo-Vento, 1998; Yoncheva et al., 2015). On the other hand, it cannot be ignored that the N1 component is primarily associated with orthographic processing, which involves visual analysis of words, it depends on the experience of perceiving letters, letter combinations and words (Bentin et al., 1999; Faísca et al., 2019; Maurer et al., 2005; Pleisch et al., 2019). In well-read adults, the N1 component, which is the electrophysiological correlate of visual word form area activation, may reflect the search for orthographic representations in the lexicon (Araújo et al., 2015). Experienced readers showed an increase in the amplitude of this component for words versus non-words, for familiar versus unfamiliar words, and for strings of letters versus strings of less familiar characters (Carreiras et al., 2014; Coch & Mitra, 2010; Mahé et al., 2012; Sánchez-Vincitore et al., 2017; Xue et al., 2019). That is, this component differentiates various orthographic stimuli and is one of the early markers of word recognition. It should be noted that in all the aforementioned studies showing differences in amplitude of the N1 component, tasks requiring a decision in relation to the presented stimuli were used. However, neither in the orthographic decision task nor in the reading task did we find any differentiation between correctly spelled words and misspelled words. This may indicate the difficulty of differentiating these correctly spelled and misspelled words even by adult native speakers, as opposed to differentiating words from pseudo-words or words from non-words. Indeed, misspelled words, unlike pseudowords, have a semantic meaning, and, unlike non-words, contain legitimate combinations of letters, which probably makes early recognition difficult. In addition, as we often encounter misspelled words in written language, these alternative forms of words generate their own mental representations which are stored simultaneously in the mental lexicon (see review by Ernestus, 2014). It can be assumed that an increase in the amplitude of N1 in the orthographic decision task is modulated by attention and reflects the depth of orthographic processing of all categories of stimuli. At the same time, given the complexity of stimulus differentiation and the probable competition of mental representations for correctly and incorrectly spelled words, recognition of the spelling of words does occur this quickly. However, a more detailed analysis of each presented word and optimization of preattentive information processing for further spelling recognition can also occur, which is reflected in the increase in the amplitude of N1 in the orthographic decision task.

In the following epoch of analysis (180-260 ms, corresponding to the P2 component), we found both the spelling effect and the interaction between the spelling effect and the task effect. At the same time, further analysis revealed differences between correctly spelled words and misspelled words only in the orthographic decision task. Moreover, although in some areas of interest (left central, central, posterior central regions) the main spelling effect was observed, in the left anterior and central areas, spelling effects were observed only in the orthographic decision task. Therefore, we supposed that the orthographic decision task predominantly modulated the spelling effect. The P2 component is associated with the extraction of orthographic and phonological features of words, that is, with the processes of decoding a grapheme into a phoneme (Barnea & Breznitz, 1998; Carreiras et al., 2005; Dambacher et al., 2006; Sánchez-Vincitore et al., 2017; Y. Wang et al., 2021). Sauseng and colleagues (Sauseng et al., 2004) reported an increased amplitude of the positive wave in the frontal regions 160 ms after the presentation of words when compared with pseudohomophones using the silent reading task followed by questions after some stimuli and the requirement to read them aloud, which could modulate differentiation processes, this increase was compatible with data obtained using the lexical decision task (Braun et al., 2009). These data are consistent with our results for the orthographic decision task on the material of words with spelling errors, which are similar to pseudohomophones. Braun and colleagues (2009) consider the P150 component (similar to the P2 component in our study) as the brain’s response to the conflict between orthographic and phonological memory representations. There is no such conflict for words, and in the case of pseudohomophones there is no orthographic representation in memory, which leads to a decrease in the amplitude of the positive wave as early as 150 ms (P150) after stimulus onset (Braun et al., 2009). This observation partly explains our finding of decreased P2 for misspelled words: although they may have orthographic representations in memory, these representations are less robust than those for words, so there may also be a conflict and the orthographic decision task predominantly modulated spelling error detection at the stage of the P2 component. However, the spelling effect can also be found in the reading task, but less strong than in the decision task. A similar spelling effect for the P2 component was observed in another study using the reading task; however, the P2 amplitude was grater for correctly than for incorrectly spelled words only in the left temporo-parieto-occipital region (Larionova & Martynova, 2022). In addition, in contrast to the present study, in the work of Larionova and Martynova (2022) a shorter stimulus presentation time of 200 ms and a larger number of stimuli were used, which could have contributed to some of the discrepancy in the results.

In addition to the spelling effect for the P2 component, a task effect was identified: the P2 amplitude in the left frontal region was greater in the orthographic decision task compared with the reading task, and in the central and parietal-temporal-occipital regions it was greater during the reading task than in the orthographic decision task. Although we did not localize the source due to the small number of electrodes, it can be assumed that these amplitude-topographic differences in the P2 component reflect processes specific to each task. Only a few neuroimaging studies have examined the effect of task on visual word recognition processes. In one of these studies comparing the lexical decision task and the silent reading task without any motor response using EEG/MEG and fMRI, Chen et al., found that task effects occurred at around 150 ms, with more activation for reading compared to a lexical decision task in precentral gyrus, as well as more activation for the lexical decision task compared to the reading task in left inferior temporal cortex and right anterior middle temporal gyrus (Chen et al., 2013). The authors attribute these differences in activation to more emphasis on early retrieval of phonological information in silent reading than in the lexical decision task (Chen et al., 2013). It is important to note that in Chen’s experiment, only words were used in the reading task, and in the lexical decision task, words and pseudowords that are clearly phonologically different from words were used, while in our study we used correctly spelled words and misspelled words that are phonologically similar to words (in two tasks). In studies using pseudohomophones, both in silent reading and in the task of lexical decision, activation was indeed noted in the region of the left inferior frontal gyrus and the left precentral gyrus (Edwards et al., 2005; Wheat et al., 2010). Therefore, we assume that phonological processes may be involved in both tasks at this stage. However, the increased P2 amplitude in the frontal regions during the orthographic decision task compared to the reading task may be associated with both general differences in attention and modulation of the task-specific process of detecting a conflict between orthography and phonology to determine the correct spelling of words. At the same time, the increase in the P2 amplitude in the posterior regions during the reading task, as well as the spelling effect found in another study in the visual word form area during reading (Larionova & Martynova, 2022) may indicate processes predominantly associated with the recognition of the spelling of the whole word, which are similar to those observed in the 90-160 ms time window in the orthographic decision task, but do not always result in the recognition of correctly spelled words.

In our study, misspelled words elicited a larger N400 component than correctly spelled words between 350 and 500 ms for both the orthographic decision task and the reading task. It is believed that a decrease in N400 amplitude may reflect easier access to lexico-semantic information (Almeida, 2021; Kutas & Federmeier, 2000; Lau et al., 2009; Meade et al., 2019). The amplitude of N400 has been shown to be larger for pseudowords than words, and this effect has been identified in various paradigms – semantic categorization (Coch & Benoit, 2015), the lexical decision task (Braun et al., 2006; Carreiras et al., 2005), the silent reading task (Bermúdez-Margaretto, Shtyrov, et al., 2020), the pronounceability judgment task (Tzeng et al., 2017). In addition, the N400 wave is associated with phonological processing and is traditionally considered in works with pseudohomophones, in which a less negative wave reflects higher activation of a particular phonological representation (Barnea & Breznitz, 1998; Briesemeister et al., 2009; Costello et al., 2021; González-Garrido et al., 2015; Y. Wang et al., 2021). The few studies using misspelled words as stimuli showed results similar to ours: N400 was more negative for misspelled words compared to correctly spelled words in the lexical decision task (González-Garrido et al., 2015), in the orthographic decision task (Heldmann et al., 2017), and in the reading task (Larionova & Martynova, 2022). According to some authors, the amplitude of N400 reflects the amount of effort required to integrate orthographic, phonological and semantic information during lexical processing (Grainger & Holcomb, 2009), and the increase in the amplitude of N400 for pseudohomophones is associated with a conflict between orthographic and phonological representations, which makes integration difficult (Briesemeister et al., 2009). We found that, regardless of the task, the N400 amplitude for misspelled words was more negative than for correctly spelled words. This is consistent with the results of other studies and suggests that spelling recognition occurred regardless of the requirements of the task. Kutas and Federmeier (2011) described in their review that the N400 includes the characteristics of both automatic and controlled processing. Our results indicate that recognition of the correct spelling of words in the time window corresponding to the N400 component occurs automatically. This probably means that spelling recognition involves lexico-semantic and phonological processes regardless of the task. It is noteworthy that we showed that the N400 amplitude was greater during passive reading than during the orthographic decision task regardless of the type of stimulus. On one hand, a decrease in the amplitude of N400 in the orthographic decision task may reflect easier access to lexico-semantic and phonological information at this stage, including due to the modulation of earlier components of evoked potentials. That is, despite the independence of the spelling recognition process, top-down modulation of the N400 amplitude can take place. On the other hand, we must take into account subtle differences between the procedures in relation to response preparation (the presence of a motor response only in the orthographic decision task, the absence of a response in the reading task), which can affect the processing of stimuli in the later stages of processing.

We have shown that the P600 amplitude is more positive in the orthographic decision task in the central and posterior areas, in contrast to the reading task, in which a negative wave was still observed in these areas in the 500-750 ms time window. At the same time, in the left anterior and middle anterior regions, P600 was more positive in the reading task than in the orthographic decision task. There was a positive wave in both tasks in these areas (Figures 1–2). It is known that the P600 component strongly depends on the task: if the task does not require an assessment of acceptability or plausibility, then this leads to a weakening of the P600 amplitude in centro-parietal areas (Kolk et al., 2003; Schacht et al., 2014). For example, in Gunter and Friederici’s study (1999), sentences with verb form violations produced N400 and P600 effects, but in a task in which participants assessed whether a word in a sentence was capitalized, the N400 and P600 components were significantly reduced or absent (Gunter & Friederici, 1999). Schacht and colleagues (Schacht et al., 2014) did not observe the parietal component of P600 in a task in which participants assessed whether a control word was present in a sentence, in contrast to the task of assessing the correctness of a sentence, so the authors concluded that P600 depends on attention to the context of sentences. The spelling task in our study required both assessment of correctness and attention to both types of presented stimuli, in contrast to the reading task, which probably explains the increase in the P600 central parietal positive wave in the spelling task.

The increase in P600 amplitude in the left anterior and middle anterior regions in the reading task compared to the orthographic decision task looks unexpected. A possible explanation for this effect may be that the P600 component in these areas in the two tasks has a different latency (Figure 2): P600 in the orthographic decision task is earlier than in the reading task, and the beginning of the P600 wave did not enter the analyzed epoch of 500-750 ms. In addition, we expected a spelling effect for the P600 component, at least in the orthographic decision task. However, we only found a trend towards significance, which was associated with the stimulus set factor and therefore was not considered. Apparently, the effects associated with the P600 component not only strongly depend on the task but also on the stimulus material.

In conclusion, we would like to note that the presence of a motor response in the orthographic decision task and its absence in the reading task means these tasks are not identical, which could affect the studied components. However, by adding a motor response to the reading task, or postponing it in the orthographic decision task, we would change the tasks themselves, which are often used in experiments, and make it difficult to compare our data with data from the literature.

## 5. Conclusions

In conclusion, the present study has demonstrated that processes associated with recognizing correct spelling of words may depend on the task. First, the orthographic decision task modulated the spelling recognition processes at 180–260 ms in contrast to the reading task. At the same time, the amplitudes of the earlier P1, N1 and P600 components were not sensitive to the spelling of words in any of the tasks. On the contrary, the amplitudes of the N400 component were modulated by the spelling of words regardless of the task. Thus, our results show that spelling recognition involves general lexico-semantic processes that are independent of the task, but the orthographic decision task modulates the spelling-specific processes necessary to quickly detect conflicts between orthographic and phonological representations of words in memory.

## Footnotes

1 at least González-Garrido et al., 2014, 2015; Larionova & Martynova 2022; Taha & Khateb, 2013 used words with real errors as stimuli

## Data availability statement

The datasets generated for this study are available on request to the corresponding author.

## CRediT authorship

Ekaterina Larionova: Conceptualization, Methodology, Investigation, Data Curation, Formal analysis, Visualization, Writing - Original Draft Preparation. Zhanna Garakh: Investigation, Writing - Review & Editing. Olga Martynova: Conceptualization, Supervision, Writing - Review & Editing.

